# Butyrate prevents visceral adipose tissue inflammation and metabolic alterations in a mouse model of Friedreich’s ataxia

**DOI:** 10.1101/2023.04.06.535845

**Authors:** Riccardo Turchi, Francesca Sciarretta, Marta Tiberi, Matteo Audano, Silvia Pedretti, Concetta Panebianco, Valentina Nesci, Valerio Pazienza, Alberto Ferri, Simone Carotti, Valerio Chiurchiù, Nico Mitro, Daniele Lettieri-Barbato, Katia Aquilano

**Author notes:** Correspondence: Katia Aquilano (lead contact),; Daniele Lettieri-Barbato. Co-Last authors (Daniele Lettieri-Barbato, Katia Aquilano).

## Abstract

Friedreich’s ataxia (FA) is a genetic neurodegenerative disease caused by mutation in *FXN* gene encoding for the mitochondrial protein frataxin (FXN). Patients with FA display an increased risk of developing diabetes that may aggravate disease prognosis. Recent studies have indicated that in addition to increased visceral adiposity, FA patients undergo a low-grade inflammatory state. The expansion of white adipose tissue (WAT) plays a fundamental role in the development of type 2 diabetes as it becomes insulin-resistant and a source of inflammatory molecules (adipokines). In this work, we have characterized visceral WAT (vWAT) at metabolic and immunological level in a murine FA model (KIKO) to test whether dysfunction of vWAT could be involved in FA metabolic complications. Through RNAseq analyses we found an alteration of inflammatory, angiogenesis and fibrosis genes in vWAT of KIKO mice. We also found other diabetes-related hallmarks such as increased lipid droplet size, immune cell infiltration and increased expression of pro-inflammatory cytokines. In addition, by targeted metabolomics we disclosed a raise in lactate production, an event usually associated with obesity and diabetes and that triggers activation of vWAT resident macrophages. To reproduce an *in vitro* model of FA, we downregulated FXN protein in cultured white adipocytes and recapitulated the diabetes-like features observed in vWAT. Real time monitoring of adipocyte metabolism evidenced metabolic rewiring towards glycolysis according to increased lactate production. Analysis of fecal samples revealed a reduction of some butyrate-producing bacteria in KIKO mice. As this microbiota derived short-chain fatty was demonstrated to exert anti-diabetic function, we fed KIKO mice with a butyrate-enriched diet for 16 weeks. This dietary approach limited vWAT alterations and mitigated other diabetes-like signatures both in *in vitro* and *in vivo* models.

In conclusion, this study identified vWAT as an important player in the onset of metabolic complications typical of FA and suggests butyrate as safe and promising adjuvant tool to treat metabolic complications in FA.

## INTRODUCTION

Friedreich’s ataxia (FA) is a rare neurodegenerative disease caused by the expansion of intronic trinucleotide repeat GAA from 8-33 repeats to >90 repeats in the *FXN* gene encoding frataxin protein (FXN). FXN resides in mitochondria and regulates mitochondrial iron transport and respiration (Llorens et al., 2019). Apart from manifesting neurodegenerative signatures, FA patients are at higher risk to developing type 2 diabetes (T2D) and cardiomyopathy than general population, and these concur to aggravate the prognosis (extensively reviewed in (Tamarit et al., 2016). It is now well ascertained that lipid metabolism is altered at systemic and cellular level in FA. FA patients show accumulation of lipid droplets (LD) in fibroblasts (Coppola et al., 2009). LD accumulation has been also observed in FXN deficient cultured rat cardiomyocytes and iPSC-derived FA cardiomyocytes (Obis et al., 2014;Li et al., 2019) as well as in heart and brown adipose tissue of FA mouse models (Puccio et al., 2001;Stram et al., 2017;Turchi et al., 2020). Hepatic accumulation of fat (steatosis) in mice with a liver-specific FXN ablation (Martelli et al., 2012) and altered lipid metabolism associated with increased LD in glial cells of the drosophila FA model (Navarro et al., 2010) have been also observed.

White adipose tissue (WAT) is the tissue with the highest capacity to accumulate fats within LD and release them to other tissues in response to increased energetic demands. In addition to being a storage depot, WAT is a high active major endocrine organ impacting the metabolic function of several tissues and overall body metabolic homeostasis (Luo and Liu, 2016). Subcutaneous WAT (sWAT) is mainly involved in the buffering of circulating free fatty acids and triglycerides, thus exerting a protective function against systemic lipotoxicity and pathological accumulation of visceral WAT (vWAT) (Frayn, 2002;Abate and Chandalia, 2012). vWAT is present mainly in the abdominal region, and its expansion largely contributes to the onset of systemic low-grade inflammatory states that are at centre stage of insulin resistance and T2D development (Burhans et al., 2018). In addition to adipocytes, vWAT contains a great number of stromal vascular cells (SVCs) including endothelial cells, preadipocytes, and immune cells. Among the immune cells, macrophages play an important role in vWAT inflammation. Indeed, concomitant to pathological expansion, the ratio between the pro-inflammatory M1 macrophages and anti-inflammatory M2 macrophages increases, thus eliciting a major production of proinflammatory cytokines (e.g., TNF-alpha) (Lu et al., 2019). Several evidence indicate that impaired oxidative function of mitochondria in vWAT adipocytes may be causally involved in its expansion and development of low-grade inflammation, insulin resistance and T2D. Of note, the amphibolic organelles mitochondria take centre stage in maintaining metabolic homeostasis in white adipocytes because of their involvement in fatty acid synthesis and esterification as well as lipid oxidation (Boudina and Graham, 2014;Barquissau et al., 2016). Notably, increased visceral adiposity along with systemic chronic, low-grade inflammatory state was observed in FA patients (Cnop et al., 2013;Nachun et al., 2018). Also, obesogenic diet, that inevitably leads to WAT expansion, was demonstrated to exacerbate the metabolic dysfunctions caused by FXN deficiency in mice, indicating a role for FXN in the maintenance of WAT function (Pomplun et al., 2007).

Efficient therapeutics to definitively cure FA have not been identified yet. Supplementation with the short-chain fatty acid butyrate ameliorates body metabolism and increases insulin sensitivity (Gao et al., 2009;Roelofsen et al., 2010;Khan and Jena, 2016). Moreover, butyrate ameliorates oxidative function of mitochondria and increases lipolysis in vWAT, thus limiting its expansion and restoring plasma leptin levels in diabetic mice (Jia et al., 2017;Pelgrim et al., 2017).

In this study we evaluated whether dysfunction of vWAT could be operative in FA and butyrate could be used as a safe dietary supplement to counteract the T2D-like features of WAT in FA.

## RESULTS

### WAT of KIKO mice show alterations typical of type 2 diabetes

We previously showed that frataxin knock-in/knockout (KIKO) mice undergo weight gain starting at 8-months of age that was accompanied by an increase in circulating levels of leptin (Turchi et al., 2020), suggesting that WAT dysfunction occurs in this mouse model. Hence, we performed bulk RNAseq analysis of the main WAT depot involved in T2D development i.e., vWAT. We found 97 differentially expressed genes (FC>1.5; FC<0.5; FDR<0.05; 41 up-, 56 down-regulated) between WT and KIKO mice (**Fig. 1A**).

**Figure 1.**
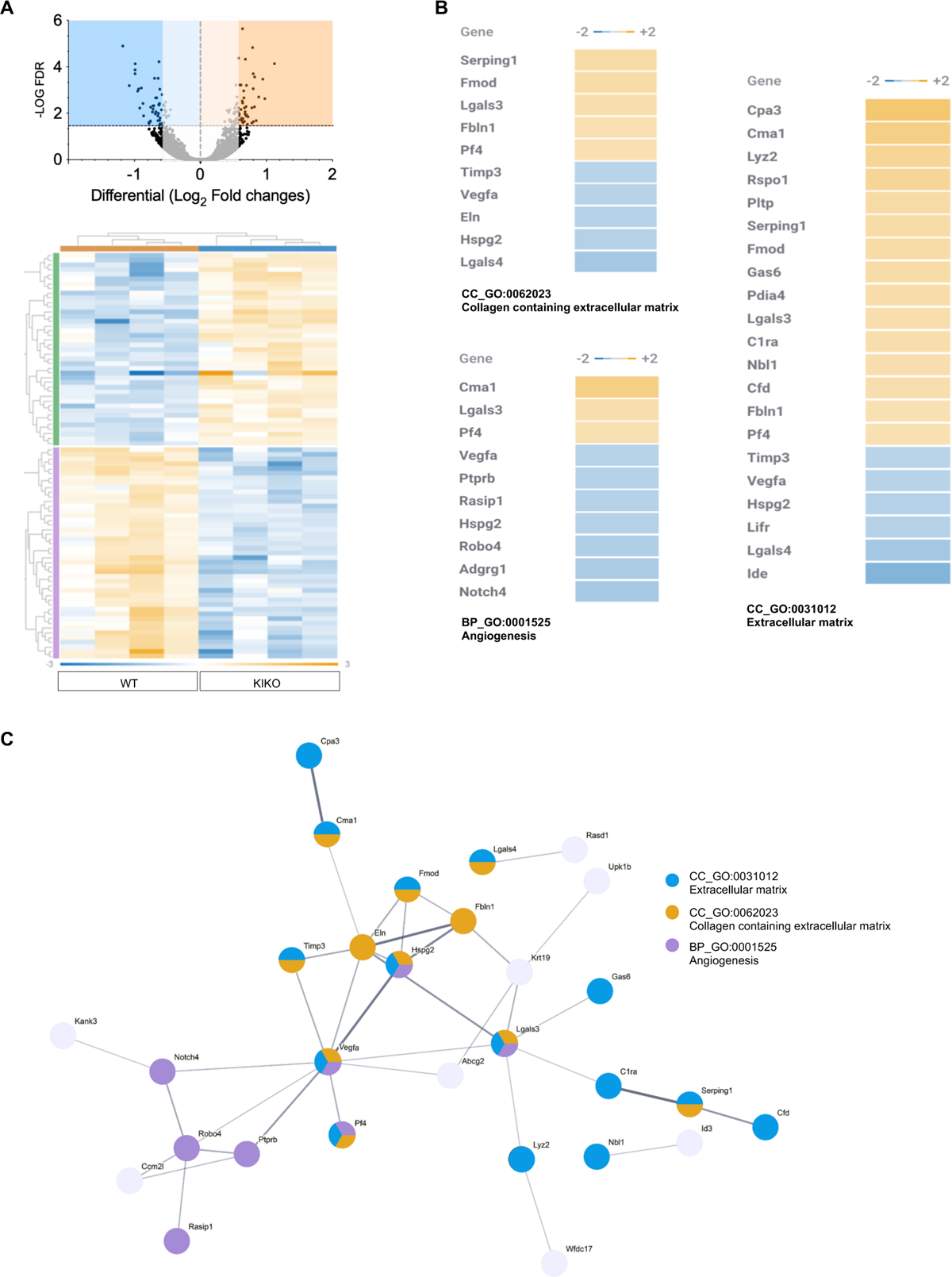
Alteration of gene expression in vWAT of KIKO mice. **A**) Volcano-plot representing differentially expressed genes between vWAT of WT and KIKO mice (upper panel). Heatmap representing the hierarchical clustering of significantly modulated genes (n=4 mice/group; fold change>1.5, <0.5; FDR<0.05) (bottom panel). **B**) Functional enrichment analysis of modulated genes. **C**) Protein-protein interaction network of modulated genes obtained by STRING platform. Nodes were coloured according to the enriched GO term.

Functional enrichment analysis of the differentially expressed genes revealed angiogenesis and extracellular matrix as the modulated biological processes and cellular component, respectively (**Fig. 1B**). Analysis of the cellular components also evidenced an enrichment of genes pertaining to collagen-containing extracellular matrix (**Fig. 1B**). In the GO term extracellular matrix, the presence of genes typically expressed and secreted by macrophages and mast cells were found. These include i) the macrophage-specific Lyz2, involved in the onset of local WAT inflammation (Latorre et al., 2021); ii) the mast cell-specific Carboxypeptidase A3 (Cpa3) and Chimerase 1 (Cma1), involved in the digestion of extracellular matrix and fibrosis development (Dell’Italia et al., 2018;Owens et al., 2019). Overall data indicated a vWAT rearrangement towards hypovascularization, inflammation and fibrosis. To find a causal relationship between differentially expressed genes, we built a protein-protein interaction network using the free STRING platform (https://string-db.org/). The network depicted in **Fig. 1C** highlights that some of the differentially expressed genes create an interconnected network with VEGFA representing a central hub. VEGFA is a growth factor inducing proliferation and migration of vascular endothelial cells and is essential for both physiological and pathological angiogenesis. RNAseq data were validated through qPCR analysis. As reported in **Fig. 2A**, *Vegfa* mRNA levels resulted downregulated along with *Rasip1* - a protein essential for the correct assembly and angiogenic migration of endothelial cells (Koo et al., 2016) - and *Robo4* - an endothelial receptor that is involved in the maintenance of endothelial barrier organization and function (Dai et al., 2019).

**Figure 2.**
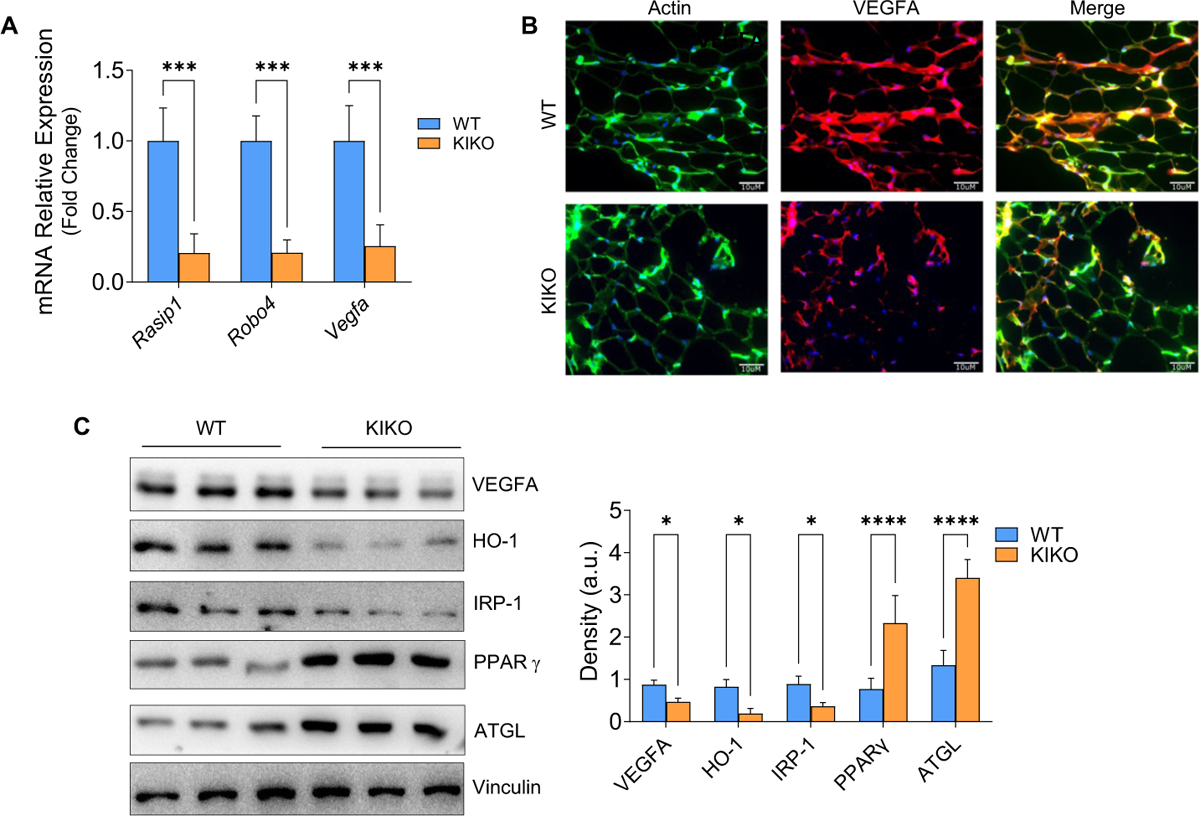
vWAT of KIKO mice show alteration of angiogenesis and lipid metabolism. **A**) RT-qPCR analysis of angiogenic genes. Data are expressed as mean ± SD (n = 4 mice/group; *** p<0.001); **B**) Representative immunofluorescence analysis of VEGFA (red). Phalloidin was used to stain F-actin (green) and Hoechst 33342 (blue) to visualize nuclei (Magnification 200×; scale bar 10 μm). **C**) Western blot analysis of proteins related to blood vessel endothelial cell proliferation (VEGFA, HO-1 and IRP-1) and lipid metabolism (ATGL, PPARγ). Vinculin was used as loading control. Density of immunoreactive bands were normalized with respect to loading control. Data are expressed as mean ± SD (n = 3 mice/group; * p<0.05, **** p<0.0001).

Immunofluorescence analysis and western blot analysis of VEGFA confirmed its downregulation in KIKO mice also at protein level (**Fig. 2B, C**). The decrease of the protein content of the downstream effectors HO-1 and IRP-1 corroborated the overall downregulation of VEGFA pathway in vWAT of KIKO mice (**Fig. 2C**). We previously demonstrated that FXN deficiency causes lipid accumulation in brown adipose tissue (Turchi et al., 2020). Western blot analysis of hallmarks of lipid loading such as the adipose triglyceride lipase (ATGL) and the transcription factor and activator of lipid synthesis PPARγ, indicated that lipid accumulation also occurs in vWAT (**Fig. 2C**).

Hypovascularization and vWAT expansion can lead to lowering of oxygen availability, decrease of mitochondrial respiration and metabolic reprogramming towards glycolysis and lactate production. Notably, increased levels of lactate in WAT are typical of T2D conditions (Feng et al., 2022;Lin et al., 2022). In order to pinpoint possible metabolic alterations of WAT, we isolated SVCs, and pre-adipocytes were induced to differentiate. Analysis of lactate concentration in culture medium showed that mature adipocytes of KIKO mice had a higher rate of lactate production than WT mice (**Fig. 3A**). Lactate was previously regarded as the waste product of glycolysis. Recently, it has emerged that lactate serves as a danger signal that promotes polarization of resident macrophages towards a pro-inflammatory M1-like state in the context of obesity (Feng et al., 2022;Lin et al., 2022). Based on this evidence and our RNAseq results, we determined the leukocyte abundance in SVCs isolated from vWAT. After isolation of CD45^+^ cells by magnetic cell sorting, we found that leukocyte content was increased in vWAT of KIKO mice (**Fig. 3B**). Next, we evaluated the presence of macrophage infiltrates by immunofluorescence analyses of vWAT sections. We used differentiation cluster 68 (CD68) as a marker for M1-like macrophages. vWAT of KIKO mice displayed a higher number of CD68^+^cells per adipocytes when compared to WAT of WT mice (**Fig. 3C**). Moreover, some crown-like structures (consisting in several macrophages surrounding a single adipocyte) typical of inflamed WAT were observable in KIKO mice (**Fig. 3C**).

**Figure 3.**
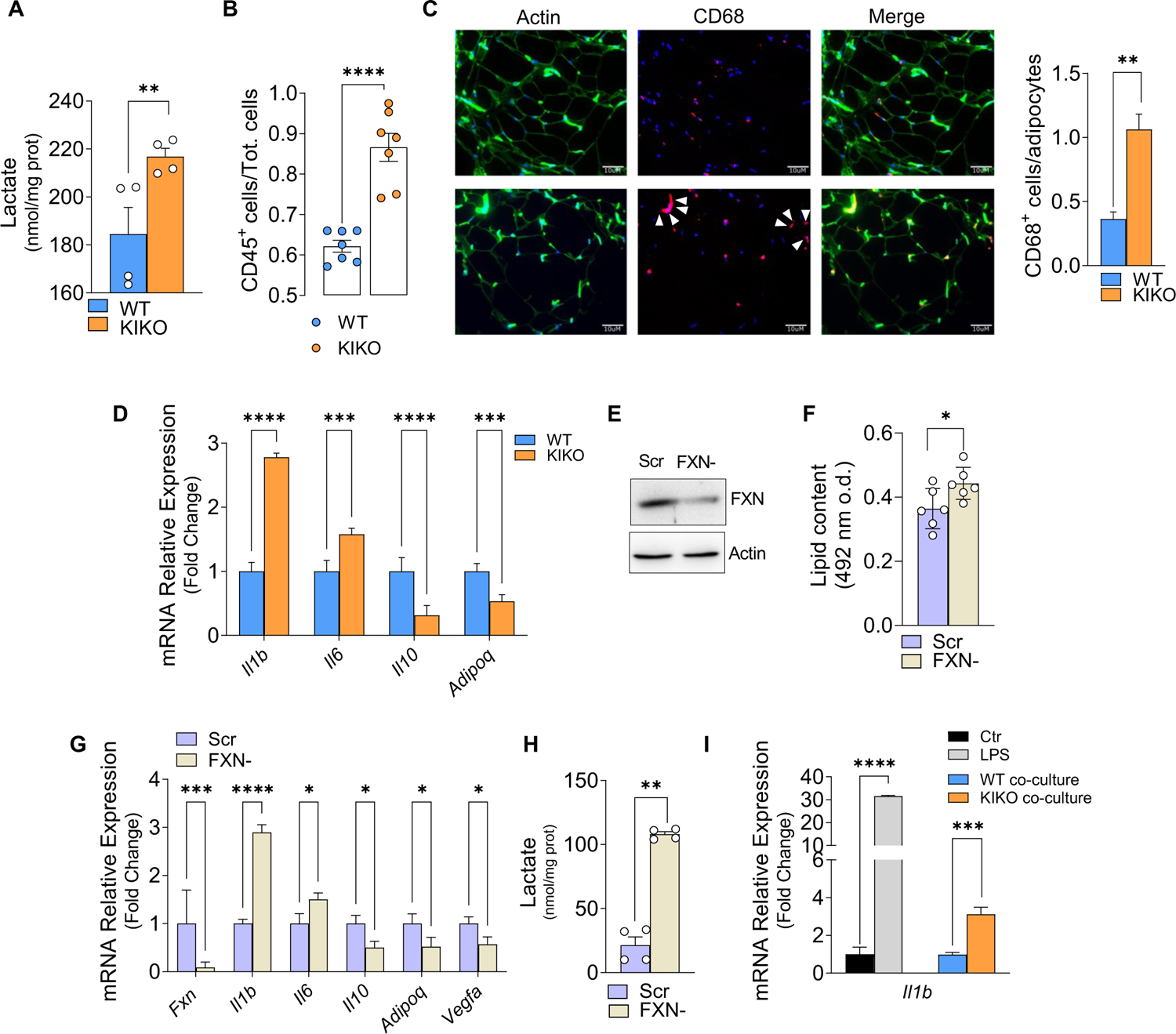
vWAT of KIKO mice show signs of macrophage infiltration and inflammation. **A**) Spectrophotometric analysis of lactate released in culture medium of vWAT adipocytes differentiated from SVCs. Data are expressed as mean ± SD (n=4 mice/group; **p<0.01). **B**) Immune cells (CD45^+^) in stromal vascular fractions obtained by magnetic cell sorting. Data are expressed as mean ± SD (n=9 mice/group; ****p<0.001). **C**) Representative immunofluorescence analysis of CD68^+^ M1 macrophage infiltrates (red). Phalloidin was used to stain F-actin (green) and Hoechst 33342 (blue) to visualize nuclei (Magnification 200×; scale bar 10 μm). White arrows indicate the presence of crown-like structures around adipocytes. Quantification of macrophage number per adipocytes is reported (*right panel*). Data are expressed as mean ± SD (n=4 mice/group; **p<0.01). **D**) RT-qPCR analysis of inflammatory genes. Data are expressed as mean ± SD (n=4 mice/group; ***p<0.001, ****p<0.0001). **E-G**) Murine 3T3-L1 adipocytes were transfected with a shRNA against FXN (FXN-) or with a Scr shRNA. Western blotting analysis of FXN and actin (loading control). Immunoblot reported is representative of three independent experiments giving similar results (E). Intracellular lipid content determined by spectrophotometric measurement of eluted Oil Red O. Data are expressed mean ± SD (n=6; *p<0.05) (F). qPCR analysis of *Fxn,* inflammatory *(Il1b, Il6, Il10, Adipoq)* and *Vegfa* mRNAs. Data are expressed as mean ± SD (n=4; *p<0.05, ***p<0.0001, ****p<0.0001) (G). Spectrophotometric analysis of lactate released in culture medium. Data are expressed as mean ± SD (n=4, **p<0.01) (H). **I**) qPCR analysis of *Il1b* mRNA in RAW264.7 macrophages co-cultured with adipocytes from WT or KIKO mice. LPS treatment was used as positive control. Data are expressed as mean ± SD (n=3; ***p<0.001, ****p<0.0001).

Thus, we investigated mRNA level of key factors involved in the regulation of the inflammatory response. vWAT of KIKO mice showed a higher expression of the pro-inflammatory *Il1b* and *Il6* genes compared to WT mice. Consistent with the hypothesis of increased inflammatory processes within vWAT of KIKO mice, anti-inflammatory *Il10* was found downregulated (**Fig. 3D**). These data were confirmed by analysing the expression of *Adipoq* gene encoding for Adiponectin, an adipokine secreted by adipocytes that inhibits macrophage-mediated inflammation (Huang et al., 2008). Interestingly, *Adipoq* mRNA expression resulted significantly decreased in vWAT of KIKO mice (**Fig. 3D**), corroborating the finding that adiponectin downregulation contributes to promoting low-grade inflammation in T2D (Fang and Judd, 2018;Funcke and Scherer, 2019).

To better elucidate the role of adipocytes in the observed macrophage recruitment and vWAT inflammation, we downregulated FXN in a cellular model of murine (3T3-L1) white adipocytes. In line with the results obtained in vWAT, FXN deficiency caused lipid accumulation (**Fig. 3E-G**), VEGFA and inflammatory marker alteration (**Fig. 3G**) extracellular lactate increase (**Fig. 3H)**. Interestingly, in co-culturing conditions, 3T3-L1 adipocytes downregulating FXN were able to induce *Il1b* gene expression in RAW264.7 macrophages (**Fig. 3I)**, pointing to a central role of adipocytes in triggering macrophage activation upon FXN deficiency.

### Dietary butyrate exerts antidiabetic effects in KIKO mice by counteracting WAT dysfunctions

Among the typical events associated with low-grade inflammation, altered gut microbiota is also included (Telle-Hansen et al., 2018). Importantly, a very strict crosstalk exists between WAT and gut microbiota that synergistically contributes to maintaining body metabolic homeostasis (Backhed et al., 2004;Shanahan et al., 2017). This prompted us to evaluate whether butyrate treatment was able to limit vWAT alteration and exert anti-diabetic functions. We supplemented mice with sodium butyrate by adding it in food pellets (5 g · kg−1 · day−1 at the normal daily rate of calorie intake) starting at 4 months of age.

Metagenomic analysis of faecal samples showed an altered microbiota composition in KIKO mice (**Fig. 4A, Table I**). Deeper analysis conducted at genus level highlighted an almost total absence of butyrate-producing bacteria in gut microbiota of KIKO mice (**Fig. 4B**). Butyrate is predominantly produced by gut microbes, and it is now ascertained that the decrease of butyrate-producing bacteria inevitably leads to diminution of systemic butyrate availability (Louis and Flint, 2017). Metagenomic analysis demonstrated that butyrate supplementation was able to restore the abundance of some butyrate-producing bacteria in faecal samples of KIKO mice to levels comparable or even larger than those of WT mice (**Fig. 4C**). These data prompted us to evaluate whether butyrate treatment was able to mitigate systemic metabolic alterations in KIKO mice. The oral glucose tolerance test (OGTT) revealed that butyrate supplementation was effective in restraining glucose intolerance in KIKO mice, as a significant recovery of normal glycaemia levels was observed after 120 min from glucose administration (**Fig. 4D**). Regarding lipidaemia, we found that butyrate was effective in buffering hypertriglyceridemia but not total cholesterol levels, even though a tendency was noted (**Fig. 4E**).

**Table I.** Antibodies used immunohistochemistry, immunofluorescence and western blot analyses.

| **Antibody** | **Source** | **Identifier** |
| --- | --- | --- |
| ACTIN | Proteintech | 20536-I-AP |
| ATGL | Cell Signaling | 2439S |
| CD68 | Abcam | ab125212 |
| FXN | Santa Cruz Biotechnology | sc-25820 |
| HO-1 | Abcam | ab13240 |
| IRP-1 | Santa Cruz Biotechnology | sc-14216 |
| PKA p-substrates | Cell Signaling | cs-9621 |
| PLIN1 | Santa Cruz Biotechnology | sc-67164 |
| PPAR-γ | Santa Cruz Biotechnology | sc-7196 |
| S100A8 | Invitrogen | PA5-86063 |
| SOD1 | Abcam | ab51254 |
| VEGFA | Santa Cruz Biotechnology | sc-7269 |
| VINCULIN | Invitrogen | MA5-11690 |

**Figure 4.**
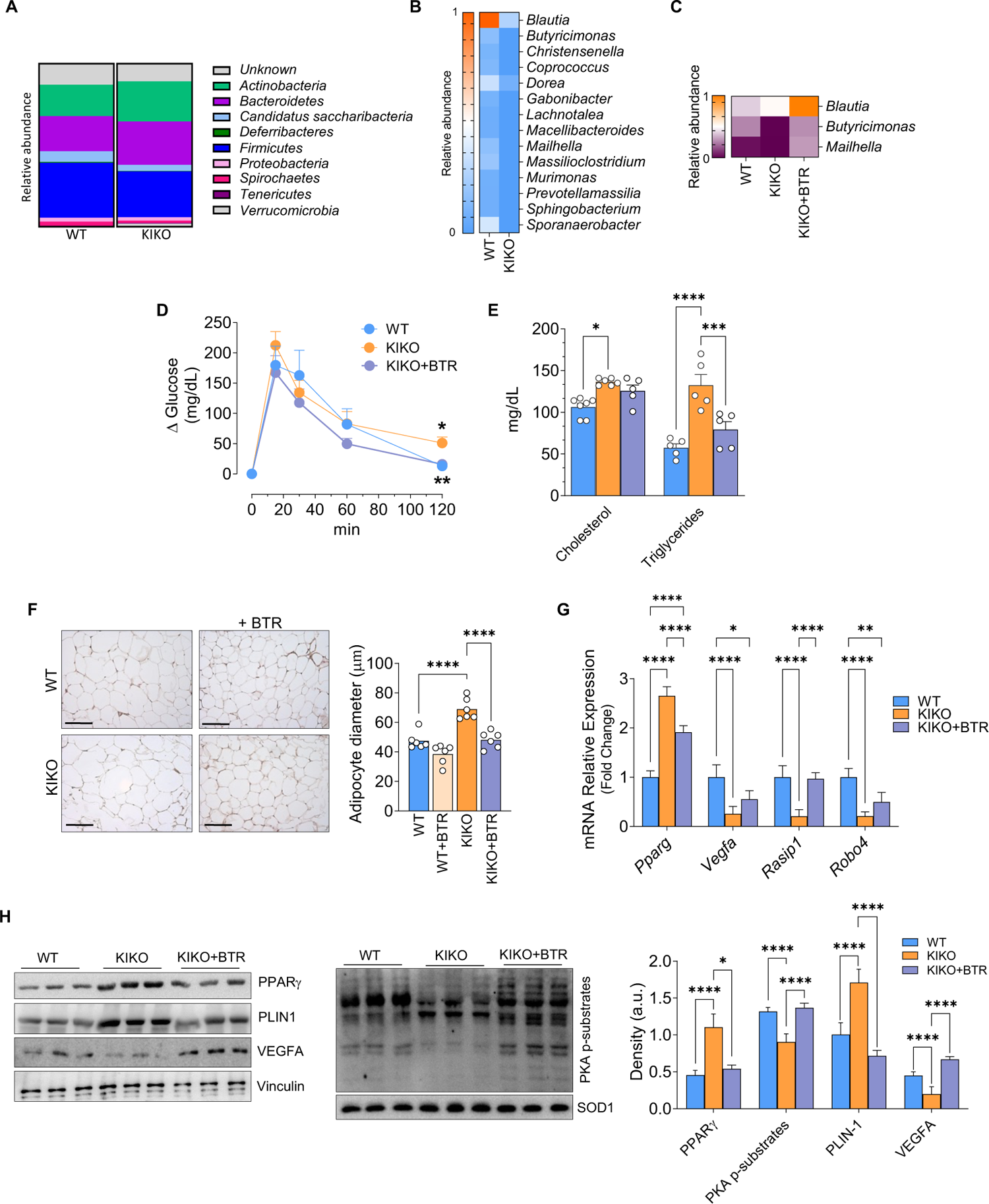
Butyrate supplementation counteracts development of T2D-like symptoms in KIKO mice. **A, B**) Relative abundance of microbiota phyla (SILVA database analysis) (A) and heatmap representation of butyrate-producing bacteria (genera in terms of relative abundance) (B) in faecal samples of WT and KIKO mice (n=12 mice/group). **C-H)** Mice were fed with normal diet or with diet supplemented with butyrate (+BTR) for 16 weeks. Heatmap representation of butyrate-producing bacteria in faecal samples whose relative abundance is modulated by butyrate in KIKO mice (n=6 mice/group) (C). Oral glucose tolerance test (OGTT). Data are expressed as mean ± SD (n=6 mice/group; *p<0.05 vs WT mice; **p<0.01 vs butyrate-untreated KIKO mice) (D). Plasma triglycerides and cholesterol levels. Data are expressed as mean ± SD (n=6 mice/group; *p<0.01, ***p<0.001, ****p<0.001) (E). Representative histology images of vWAT from mice fed with normal diet or treated with butyrate (+BTR) after staining with H&E (F, left panel). Adipocyte size was represented as lipid droplet diameters (F, right panel). Data are expressed as mean ±SD (n=6 mice/group, ****p<0.0001). RT-qPCR analysis of *Pparg* and angiogenic mRNAs. Data are expressed as mean ±SD (n = 3 mice/group; *p<0.05; ** p<0.01; **** p<0.0001) (G). Western blot analysis of proteins involved in lipid metabolism and VEGFA. SOD1 and Vinculin were used as loading control. Density of immunoreactive bands were normalized with respect to related loading control. Data are expressed as mean ± SD (n=3 mice/group; *p<0.01, ****p<0.001).

Histochemical evaluation displayed an increase in adipocyte size in vWAT of KIKO compared to WT mice, confirming vWAT expansion (**Fig. 4F**). Notably, butyrate treatment was able to reduce the diameter of adipocytes in vWAT of KIKO mice, with vWAT adipocytes of butyrate-treated KIKO mice reaching a size comparable to that observed in vWAT adipocytes of untreated WT mice (**Fig. 4F**). These results led us to evaluate whether vWAT dysfunction could be recovered by butyrate treatment. Western blot and qPCR analyses showed that markers of lipid accumulation (PPARγ, PLIN-1) and hypovascularization (VEGFA, Rasip, Roboa4) came back to control values upon butyrate treatment (**Fig. 4G, H**). Accumulation of lipids depends on increased PPARγ-mediated lipogenesis and inhibition of the hormone-sensitive lipolytic cascade activated by protein kinase A (PKA) (Luo and Liu, 2016). By Western blot analysis we observed a decrease of the level of PKA-phospho-substrates, and butyrate was able to revert this event (**Fig. 4H**). Overall, these results underline the beneficial effects of butyrate in maintaining lipid homeostasis and vascularization in vWAT of KIKO mice.

Immunohistochemical analyses also proved that, along with the recovery of VEGFA levels, collagen content and immune cell infiltrates were reduced upon butyrate treatment (**Fig. 5A**). In parallel, butyrate restrained the upregulation of the pro-inflammatory *Il1b* and *Cox*2 genes and restored the mRNA level of the anti-inflammatory cytokine *Il10* (**Fig. 5B**). Lactate also represents a metabolic mediator that shapes immune cell dynamics within WAT depots (Caslin et al., 2021). Accordingly, we sought to deeply characterize innate and adaptive immune cell dynamics by investigating potential recruitment of other immune cell populations in vWAT and whether butyrate was able to exert an impact on this. To this end, we isolated SVCs of vWAT, and the single cell suspension was analysed by high dimensional flow cytometry. As expected, a higher percentage of total CD45^+^ leukocytes was observed in SVCs of KIKO compared to WT mice (**Fig. 5C**). We then applied a consequential gating strategy to identify the percentages of the different cell subsets of leukocytes, *i.e.,* macrophages (CD11b^+^F4/80^+^CD64^+^ cells), neutrophils (Ly6G^+^ Ly6C^-^ cells), T cells (CD3^+^ cells), natural killer (NK) (NK1^+^ cells) and B cells (CD19^+^ cells). Interestingly, vWAT of KIKO mice showed a higher percentage of macrophages and neutrophils compared to WT mice (**Fig. 5C**), corroborating our previous findings by immunohistochemical analyses (**Fig. 3C, 5A**). By contrast percentages of T and B lymphocytes as well as NK cells was not changed (**Fig. 5C**), suggesting that NK cells and cells of the adaptive immunity do not contribute to the inflammation of vWAT in KIKO mice. Notably, butyrate was able to prevent leukocyte infiltration, and this was due to reduction of both macrophages and neutrophils (**Fig. 5C**).

**Figure 5.**
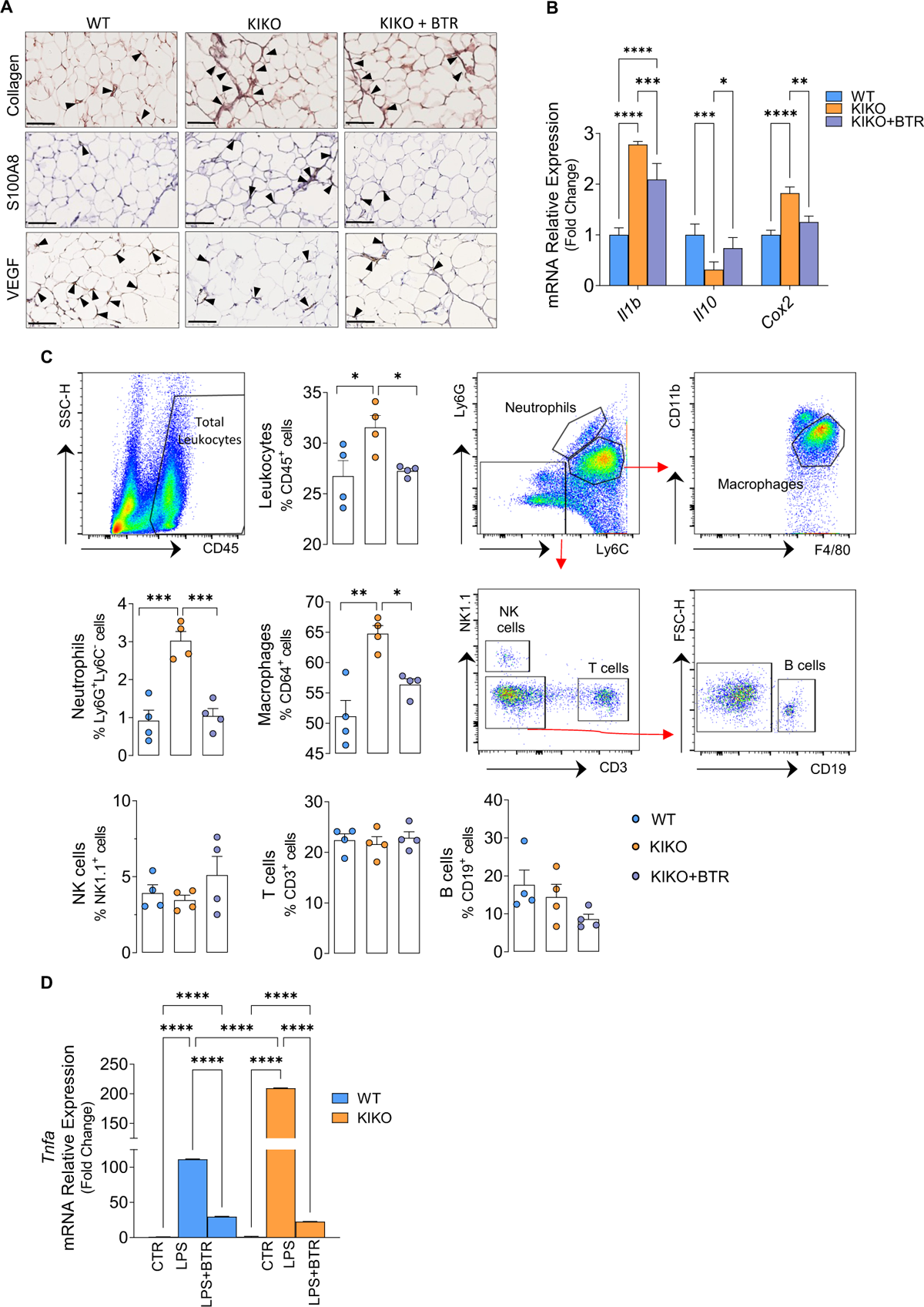
Butyrate treatment prevents the onset of inflammation in vWAT of KIKO mice. Mice were fed with normal diet or with diet supplemented with butyrate (+BTR) for 16 weeks. **A)** Representative histology images of vWAT to detect collagen (masson trichrome staining), VEGFA (staining with VEGFA antibody) or immune cell infiltrates (staining with S100a8 antibody). Arrowheads indicate fibrotic septa, VEGFA positive staining or inflammatory infiltrates with scattered cells (Magnification 200×; scale bar 10 μm). **B**) RT-qPCR analysis of inflammatory genes. Data are expressed as mean ± SD (n = 3 mice/group; *p<0.05, ** p<0.01, ***p<0.001, **** p<0.0001). **C**) SVCs were isolated and analysed through high dimensional flow cytometry by using specific antibodies to detect total leukocytes, neutrophils, macrophages, T cells, NK cells and B cells. Gating strategies to detect the immune cell subpopulations are illustrated. Data are expressed as mean percentage of positive cells ± SD (n=4 mice/group; *p<0.05, **p<0.01, ***p<0.001). **D**) RT-qPCR analysis of *Tnfa* mRNA in BMDM of WT and KIKO treated with LPS (500 ng/ml, 16h) alone or in combination with butyrate (500 nM). Data are expressed as mean SD (n=3; ****p<0.00001).

Immune cells have higher capacity to produce inflammatory cytokines than adipocytes; hence, once recruited in vWAT by adipocyte derived signals, immune cells could enhance the production of inflammatory mediators. Hence, we questioned whether FXN deficiency could also influence the inflammatory response in immune cells and butyrate to restrain this event. To this end, we isolated bone marrow derived macrophages (BMDMs) from WT and KIKO mice. BMDMs were treated with lipopolysaccharide (LPS) to reproduce an inflammatory insult. Up-regulation of the expression of the *Tnfa* gene was obtained after LPS stimulation in both WT and KIKO BMDMs (**Fig. 5D**); however, even though FXN deficiency did not influence basal *Tnfa* expression, upon LPS treatment *Tnfa* up-regulation was more marked in KIKO than WT cells. Notably, butyrate was able to significantly buffer the LPS mediated inflammatory challenge (**Fig. 5D**), suggesting that this molecule has an anti-inflammatory action also in myeloid cells and likely at systemic level.

To have a comprehensive view of the effects of butyrate on WAT metabolism, we then performed targeted metabolomic analyses of several metabolites (about 100) pertaining to glycolysis, pentose phosphate pathway, urea and Krebs cycle and other metabolites such as carnitines, nucleotides, amino acids and catecholamines (**Table II**). Only few metabolites were significantly affected in vWAT of KIKO mice (i.e., lactate, oxalacetate and succinate) (**Fig. 6A**). Among these, only lactate underwent significant reduction upon butyrate treatment, according to data from *in vitro* experiments. We then performed XF Seahorse real-time monitoring of cell metabolism in SVCs isolated from vWAT of KIKO mice to understand whether lactate hyperproduction observed with metabolomic analyses was associated with augmented glycolytic rate, and whether butyrate treatment was able to mitigate such phenomenon *in vivo*. Lactate production of KIKO SVCs was attenuated following 16 weeks treatment with butyrate (**Fig. 6B**). Expectedly, in SVCs of KIKO mice we found a significant decrease of spare respiratory capacity, which is a measure of the ability of mitochondria to respond to increased energy demand (**Fig. 6C**). The monitoring of acidification rate (ECAR), after addition of saturating amount of glucose, revealed that KIKO SVCs have higher glycolytic rate than WT adipocytes (**Fig. 6C**). The addition of the mitochondrial respiration inhibitor oligomycin revealed that also the maximum glycolytic capacity was higher in KIKO than WT adipocytes (**Fig. 6C**). Notably, butyrate was able to recover mitochondrial respiration capacity and to reduce glycolytic metabolism in KIKO SVCs (**Fig. 6C**). We also tested the effects of butyrate on lactate production of SVCs after induction of adipocyte differentiation. As reported in **Fig. 6D**, treatment with butyrate lowered the concentration of lactate in culture medium of KIKO adipocytes. These results point to the ability of the *in vivo* treatment with butyrate to shift cellular metabolism from glycolysis to mitochondrial respiration, thus avoiding the release of anti-lipolytic and pro-inflammatory lactate and cytokines in WAT.

**Table II.**
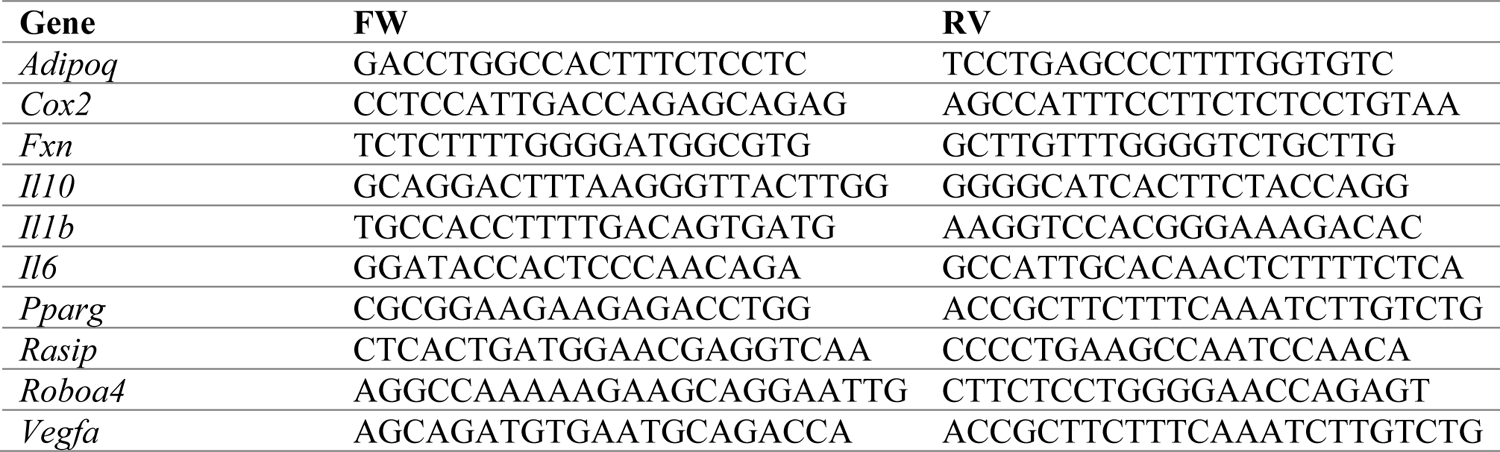
Primers used for RT-qPCR analysis.

**Figure 6.**
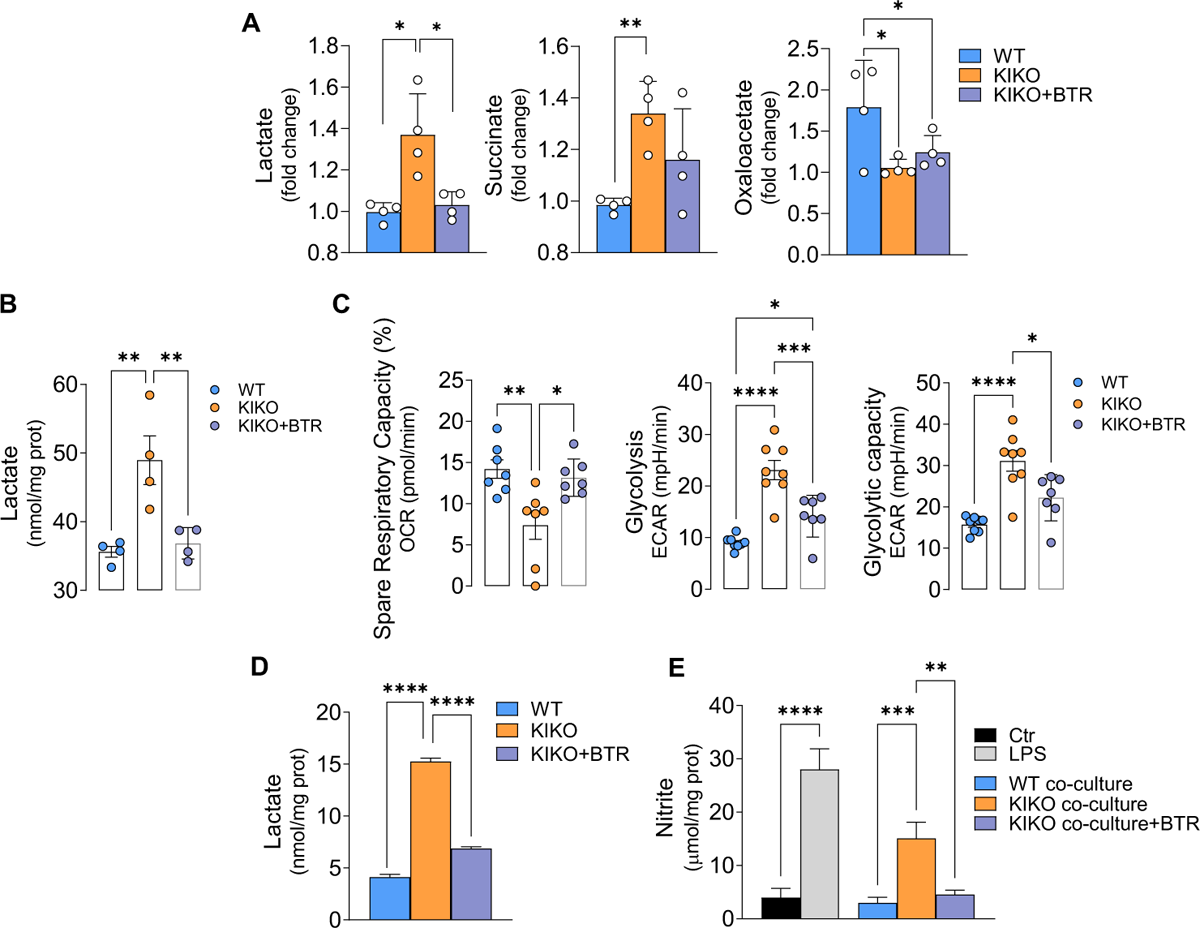
Butyrate treatment improves adipocyte metabolism in KIKO mice. Mice were fed with normal diet or with diet supplemented with butyrate (+BTR) for 16 weeks. **A**) Targeted metabolomic analysis of vWAT. Data are expressed as mean ± SD (n=4 mice/group; *p<0.05, **p<0.01). **B)** Spectrophotometric analysis of lactate released in culture medium of vWAT SVCs. Data are expressed as mean ± SD (n=4 mice/group; **p<0.01). **C**) Real-time measurement of Spare Respiratory capacity, glycolysis and glycolytic capacity. ECAR: ExtraCellular Acidification Capacity. Data are expressed as mean ± SD (at least n=6 each group; * p<0.05; **p<0.01; ***p<0.001; ****p<0.0001). D) Spectrophotometric analysis of lactate released in culture medium of differentiated adipocytes from vWAT and treated with butyrate (500 nM, 16 h). Data are expressed as mean ± SD (n=3; ****p<0.00001). **D**) Nitrite concentration in culture medium of RAW264.7 macrophages co-cultured with adipocytes from WT or KIKO mice and treated with BTR (500 nM, 16h). LPS treatment (500 ng/ml, 16h) of RAW264.7 macrophages was used as positive control. Data are expressed as mean ± SD (n=3; ***p<0.001, ****p<0.0001).

## DISCUSSION

In this study, we demonstrated that FXN deficiency leads to vWAT dysfunction, which consists of the expansion of adipocyte size, hypovascularization, production of pro-inflammatory adipokines, immune cell recruitment, and fibrosis. These findings recapitulate what is observed under T2D conditions in vWAT.

We previously demonstrated that FXN deficiency causes lipid accumulation and lower thermogenic capacity in brown adipocytes (Turchi et al., 2020), indicating a possible involvement of decreased anti-diabetic activity of BAT in the onset of metabolic complications observed in FA patients. As in BAT, in vWAT of our FA mouse model we found altered lipid metabolism due to increased expression of lipogenic markers and impaired lipolytic activity. This result is in line with the findings that individuals with FA and insulin resistance exhibit an altered adipose tissue distribution, with increased fat deposition in the visceral region (Cnop et al., 2012). It appears that vWAT is more affected than BAT in FA mice, inasmuch as besides fat accumulation, we found altered expression of genes related to angiogenesis, including VEGFA, the master regulator of this process also in vWAT (Corvera and Gealekman, 2014;Crewe et al., 2017). Insufficient angiogenic potential is a dominant contributor to the dysfunctional WAT, as hypoxic condition triggers chronic low-grade inflammation predominantly characterized by proinflammatory macrophage infiltration (Crewe et al., 2017). Intriguingly, VEGFA also exerts an anti-diabetic action by functioning as a promoter of insulin sensitivity and as an anti-inflammatory M2 macrophage attractant in WAT (Elias et al., 2013), suggesting that the VEGFA down-regulation observed in vWAT may participate in the development of insulin resistance and inflammation.

We demonstrated that FXN deficient white adipocytes are *per se* more prone to produce pro-inflammatory cytokines than normal adipocytes, arguing that they can play an active role in macrophage recruitment and activation. In vWAT of KIKO mice, we also disclosed high degree of immune cell infiltrates, mostly macrophages and neutrophils, in association with the upregulation of fibrosis markers (e.g., collagen) and pro-inflammatory cytokine production. These findings recapitulate the fibrotic inflammatory phenotype typical of vWAT in T2D (Corvera and Gealekman, 2014;Li et al., 2021).

Enlarged vWAT can release hormones and other substances that may contribute to insulin resistance, cardiomyopathy, and low-grade inflammation (Freitas Lima et al., 2015;Feijoo-Bandin et al., 2020). Apart from T2D and cardiomyopathy, signs of low-grade inflammation with high levels of inflammatory cytokine expression in circulating leukocytes were found in FA patients (Nachun et al., 2018). Additionally, FXN downregulation enhanced inflammatory response in macrophages, indicating that a positive loop of inflammation occurs in vWAT of FXN deficient mice, with adipocytes favouring the infiltration of immune cells that are more responsive to inflammatory stimuli. This evidence and our results point to the role of FXN in the maintenance of overall body immune homeostasis.

A number of data have highlighted that visceral adiposity and low-grade inflammation are correlated with an increased risk of developing neurological disorders and cognitive decline (Isaac et al., 2011; de et al., 2023). Altered expression of adipokines may disrupt either brain or heart homeostasis and functions (Jia et al., 2016;Parimisetty et al., 2016;Nishida and Otsu, 2017). Based on these findings, we can hypothesize that, apart from being a central event in the setting of T2D-like symptoms, vWAT alteration contributes to aggravating neurological symptoms and promoting heart failure in FA patients.

Augmented lactate levels in adipose tissue and plasma are widely reported in obese/diabetic subjects (Krycer et al., 2020;Feng et al., 2022). Metabolically, we found that FXN-deficient adipocytes have increased glycolytic activity and consequent hyperproduction of lactate. The switch towards glycolytic metabolism may originate from defective mitochondrial respiration and/or scarce oxygen availability due to reduced angiogenesis and vascularization. Lactate is a redox molecule that, in adipose tissues, displays a wide range of biological effects both through its binding to membrane receptors and its transport and subsequent effect on intracellular metabolism (Lagarde et al., 2021). Lactate produced by adipocytes inhibits lipolysis in an autocrine/paracrine manner through inhibition of PKA (Ahmed et al., 2010). Accordingly, we observed a decrease of PKA activity in vWAT of KIKO mice. High lactate levels increase cardiovascular risk (Crawford et al., 2010;Aleksandar et al., 2016;Krycer et al., 2020) and promote macrophage recruitment and WAT inflammation (Feng et al., 2022). We found that co-culturing macrophages with FXN deficient adipocytes promotes a more efficient inflammatory response. This result suggests that, apart from the increased production of inflammatory cytokines, FXN deficient adipocytes produce higher level of lactate that could be responsible for macrophage activation.

Dysbiosis has been linked to T2D and various neurological disorders such as multiple sclerosis, and autism spectrum disorders (Finegold et al., 2010;Berer et al., 2011;Cho et al., 2012). A very strict crosstalk exists between vWAT and gut microbiota that synergistically contribute to maintaining body metabolic homeostasis. Indeed, altered gut microbial ecosystems have been associated with vWAT dysfunction (i.e., expansion, inflammation, and insulin resistance), low-grade chronic inflammation, and systemic metabolic perturbations, including T2D (Cox et al., 2015). A decrease in butyrate-producing bacteria has been causally involved in vWAT expansion and inflammation, as well as in T2D development (Qin et al., 2012;Canfora et al., 2015). To our knowledge, no attempts have been made to unravel whether the microbiota is altered in FA. Herein, we show that our FA mouse model has an altered gut microbiota composition compared to healthy mice, with a reduction in the abundance of butyrate-producing bacteria. The causes of such reduction have not been explored in the present work and the possible occurrence of gut inflammation deserves further and deeper investigation. Mitochondrial metabolism of colonocytes, by consuming O_2_, maintains the predominance of anaerobic bacteria in the gut, including butyrate-producing bacteria (Litvak et al., 2018). It can be argued that mitochondrial dysfunction, which is likely to occur in FXN-deficient colonocytes, increases oxygen availability and the proliferation of facultative anaerobic bacteria, while reducing the abundance of butyrate-producing bacteria (dysbiosis). This dysbiosis could contribute to the establishment of a T2D-like inflammatory state in vWAT.

While further research is needed to fully understand the role of the microbiota in FA, our findings suggest that alterations in the microbiota may be involved in the pathophysiology of the disease and provide a potential target for therapeutic interventions. Strategies that target the gut microbiota, such as butyrate supplementation, probiotics or prebiotics, may have potential for improving symptoms and disease progression in FA.

The decrease in butyrate-producing bacteria strongly indicates that systemic butyrate availability could be reduced in KIKO mice. Butyrate has been studied for its potential anti-inflammatory and neuroprotective effects, as well as its role in diabetes management (Bayazid et al., 2022;Xiong et al., 2022). Our data demonstrate that butyrate supplementation may be effective in counteracting vWAT dysfunction and T2D-like symptoms in the context of FA. Specifically, at the metabolic level and similarly to what was previously described in colonocytes (Donohoe et al., 2011), butyrate increased respiratory capacity of mitochondria with FXN deficiency in vWAT, while decreasing glycolytic activity and lactate production. In parallel, butyrate hindered the accumulation of fats, and this was accompanied by the maintenance of angiogenic VEGFA levels to levels comparable to those of healthy mice. As also described in other models of T2D (Xu et al., 2018;Arora and Tremaroli, 2021), butyrate supplementation ameliorated glycaemic profile in KIKO mice. Regarding inflammatory signatures, butyrate reduced leukocyte infiltration (i.e., macrophages, neutrophils), and production of pro-inflammatory cytokines. Butyrate has a wide range of pleiotropic effects and mechanisms of action. Herein, although the precise mechanisms by which butyrate acts in our FA models have not been deeply investigated in this study, it is likely that the beneficial effects of butyrate can be mediated by the inhibition of histone deacetylases, thus epigenetically modulating the expression of genes involved in inflammation and energy metabolism (Berni Canani et al., 2012). Butyrate can also have a direct action on mitochondria. For instance, being a short-chain fatty acid, butyrate can be directly funnelled into mitochondria and enhance mitochondrial respiration and fatty acid oxidation and impede lipid accumulation (Donohoe et al., 2011). However, we cannot exclude that the beneficial effects of butyrate supplementation could be also dependent on the recovery of gut butyrate-producing bacteria as disclosed in FA mice.

Overall, our results suggest that vWAT is dysfunctional and microbiota altered in FA. Butyrate supplementation prevents vWAT expansion and inflammation as well as the development of T2D-like features in FA animals. To validate these results, further analysis of the microbiota and adipokines in the faeces and plasma of FA patients is warranted. These analyses on patients and the completion of the identification of the molecular mechanisms underlying the butyrate-mediated beneficial effects will hopefully pave the way for its safe usage as an adjuvant for treating T2D-related symptoms in FA.

## MATERIALS AND METHODS

### Animals and treatments

Mouse experimentation was conducted in accordance with accepted standard of humane animal care after the approval by relevant local (Institutional Animal Care and Use Committee, Tor Vergata University) and national (Ministry of Health, licenses n° 324/218-PR and n° 210/202-PR) committees. Mice were maintained at 21.0 ± °C and 55.0 ± 5.0% relative humidity under a 12 h/12 h light/dark cycle (lights on at 6:00 AM, lights off at 6:00 PM). Food and water were given *ad libitum*. Experiments were carried out according to institutional safety procedures.

Knock-in knock-out (KIKO) mice were purchased from Jackson Laboratories (#012329). Littermate C57BL/6 mice (WT) were used as controls. Researchers were blinded to genotypes at the time of testing. Butyrate supplementation was carried out by adding sodium butyrate in food pellets (5 g · kg−1 · day−1 at the normal daily rate of calorie intake) starting at 4 months of age up to 8 months of age (16 weeks treatment).

After cervical dislocation, vWAT tissues were explanted, immediately processed or stored at −80°C.

### Bio-clinical analyses

Prior to bio-clinical analyses, mice were starved for 2 h. After blood collection, bio-clinical analyses were performed by colorimetric methods. In particular, cholesterol and triglyceride levels were measured through the automatized KeyLab analyser (BPCBioSed, Italy) using specific assay kits (BPCBioSed).

For the glucose tolerance test (OGTT), mice were subjected to fasting for 12 h, followed by oral gavage with 2 g of dextrose/kg body mass. At the indicated time points, blood was collected from the tail vein and glycaemia measured using a glucometer (Bayer Countur XT, Bayer Leverkusen, Germany).

### Cells, treatments and transfection

Murine 3T3-L1 cell line were purchased from ATCC (Manassas, VA, USA), and cultured and differentiated in adipocytes according to ATCC protocol. Oil Red O was used to detect intracellular triglycerides content as previously described (Lettieri-Barbato et al., 2018).

Murine RAW 264.7 macrophages (ATCC) were cultured in DMEM supplemented with 10% FBS and 1% P/S (Life Technologies). All cells were maintained at 37°C in a humidified incubator containing 5% CO_2_.

For gene silencing, mature 3T3-L1 adipocytes were transfected with FXN shRNA scramble shRNA (Origene, Rockville, MD, USA), using LipofectamineTM 2000 transfection reagent (ThermoFisher) according to manufacturer’s instructions. Cells were used 48 h after transfection.

Twenty-four hours after plating, RAW264.7 cells were used for the experiments such as co-culturing with white adipocytes or treatment with 100 ng/mL LPS for 16 h.

### Histochemical and immunohistochemistry analysis

vWAT was stained with H&E or Trichrome staining to visualize general morphology and collagen deposition, respectively. For morphometric analyses, individual slides were digitized using the NanoZoomer Digital Microscope (Hamamatsu, Japan), and digital images were analysed using Image-J to measure the diameter of adipocytes. Values are the means of 10 fields taken from different tissue sections per mouse. Immunohistochemical detection of VEGFA or S100-A8 was performed on 3- to 5-lm-thick sections obtained from formalin-fixed tissue embedded in paraffin. Antigen retrieval was performed with Citrate Buffer (pH 6) (Dako, Glostrup, Denmark). Immunohistochemical staining was performed with anti VEGFA or anti-S100A8 (Table I). Incubations with primary antibodies were carried out for 2 hrs. Negative controls were obtained by omitting primary antibodies. The immunohistochemical procedure was performed using the MACH4 detection system (Biocare, Concord, CA, USA) and Betazoid DAB (Biocare, Concord) as a chromogen.

### Immunofluorescence analyses

Sections of frozen vWAT were incubated with permeabilization solution (PBS/Triton X100 0.2% [v/v]), blocked for 1 h by a blocking solution (PBS/BSA 5% [v/v]), and then incubated for 18 h with CD68 or VEGFA primary antibodies (**Table I**). After washing with cold PBS, sections were incubated 1 h with AlexaFluor-488 or −568-conjugated secondary antibodies (ThermoFisher Scientific). Nuclei were stained with 10 μg/mL Hoechst 33342 (ThermoFisher Scientific). AlexaFluor-488 Phalloidin (ThermoFisher Scientific) was used to stain actin. Images were acquired using an Olympus IX-81 confocal microscope at 60× magnitude. Representative regions of interest were acquired using a digital 3× zoom. Fluorescence intensities were set for the control samples and were maintained for all samples. To evaluate macrophage infiltration, CD68^+^ cells around adipocytes were counted in each field (10 fields/sample).

### Immunoblotting

Tissues or cells were lysed in RIPA buffer (50 mM Tris-HCl, pH 8.0, 150 mM NaCl, 12 mM deoxycholic acid, 0.5% Nonidet P-40, and protease and phosphatase inhibitors). Five μg proteins were loaded on SDS-PAGE and subjected to Western blotting. Nitrocellulose membranes were incubated with primary antibodies (**Table I**) at 1:1000 dilution. Successively, membranes were incubated with the appropriate horseradish peroxidase-conjugated secondary antibodies. Immunoreactive bands were detected by a FluorChem FC3 System (Protein-Simple, San Jose, CA, USA) after incubation of the membranes with ECL Selected Western Blotting Detection Reagent (GE Healthcare, Pittsburgh, PA, USA). Densitometric analyses of the immunoreactive bands were performed by the FluorChem FC3 Analysis Software.

### Bulk RNA-sequencing and functional enrichment analysis

Total vWAT RNA was isolated using TRIzol reagent (Invitrogen, Waltham, MA, USA) and purified using the RNeasy mini kit protocol (Qiagen, Hilden, Germany) according to the manufacturer’s instructions. Isolated RNA was sequenced using an Illumina NextSeq500, and the indexed libraries were prepared from 1 μg of purified RNA with TruSeq-stranded mRNA (Illumina) Library Prep Kit according to the manufacturer’s instructions. The quality of the single-end reads was evaluated using FastQC version 0.11.5 (https://www.bioinformatics.babraham.ac.uk/projects/fastqc). All FastQC files were filtered to remove low-quality reads and adapters using Trimmomatic version 0.36 (Bolger et al., 2014). The resulting reads were mapped to the *Mus musculus* genome (GRCm38) using HISAT2 version 2.1.0 (Kim et al., 2015) using default parameters, and StringTie version 1.3.4d (Pertea et al., 2015) was applied to the BAM files obtained using HISAT2 to generate expression estimates and to quantify the transcript abundance as transcripts per kilobase per million of mapped reads. The count matrices generated by StringTie were imported in R, in which differential expression analysis was performed using Deseq2 to compare the two different conditions. Functional annotation was performed using the AnnotationDbi R library (http://bioconductor.org). Differentially expressed genes were selected with a threshold of FC > 1.5 or <0.5 (FDR < 0.05). Functional enrichment analyses were performed using Rosalind version 3.36.1.2

### RT-qPCR

Total RNA was extracted using TRI Reagent® (Sigma-Aldrich). RNA (3 μg) was retro-transcribed by using M-MLV (Promega, Madison, WI). qPCR was performed in triplicate by using validated qPCR primers (BLAST), Applied Biosystems™ *Power*™ SYBR™ Green Master Mix, and the QuantStudio3 Real-Time PCR System (ThermoFisher, Whaltam, MA, USA) as previously described (Aquilano et al., 2016). mRNA levels were normalized to actin mRNA, and the relative mRNA levels were determined through the 2^−ΔΔCt^ method. The primers used for RT-qPCR are listed in **Table II**.

### Isolation of stromal vascular cells and bone marrow derived macrophages

Stromal vascular cells (SVCs) were obtained by finely mincing vWAT with scissors, followed by incubation in high-glucose DMEM containing 0.1% collagenase II at 37°C in an orbital shaker (150 rpm, 1h). The resulting cell suspension was then filtered through a 100-μm nylon mesh to remove any remaining tissue fragments. The filtered cells were collected in a 50-mL conical tube and centrifuged at 500 × g for 5 minutes at 4°C to remove any supernatants and floating adipocytes. Pellet was resuspended in 1 mL of ACK RBC lysis buffer and incubated at room temperature for 2 min to remove any remaining red blood cells. The RBC lysis reaction was then quenched by adding 10 mL of cold wash media (high-glucose DMEM containing 5% heat-inactivated FBS, L-glutamine, and penicillin/streptomycin). The cells were then centrifuged again at 500 × g for 5 minutes at 4°C, and the resulting pellets containing the SVCs were collected.

For isolation of bone marrow derived macrophages (BMDMs), bone marrow was extracted from the limbs of 8-months-old male mice by perfusion with PBS and 1% P/S. BMDMs were plated at a density of 3 × 10^5^ cells/mL in alpha-MEM supplemented with 10% FBS, 1% P/S, and 1% GlutaMAX. Macrophage differentiation was induced by adding M-CSF (20 ng of cells/mL) in the culture medium for 5 days. Following adhesion, unattached cells were removed, and BMDM were used for the experiments.

### Magnetic cell sorting

SVCs were resuspended in 500 mL of magnetic bead buffer (MBB) consisting of PBS without calcium and magnesium, 0.5% w/v bovine serum albumin (BSA), and 2 mM ethylenediaminetetraacetic acid (EDTA). The cell suspension was then filtered through a 30-mm pre-separation filter (Miltenyi, Bergisch Gladbach, Germany) following three filter washes to remove any large particles and debris.

The resulting cell suspension was then separated at 300 × g for 5 min at 4°C and resuspended in 90 mL of MBB along with 10 mL of anti-CD45 magnetic beads-conjugated antibody (Miltenyi) for each 10^7 cell sample. The cell suspension was incubated for 15 min at 4°C, then diluted with 2 mL of MBB and centrifuged. The resulting cell pellet was resuspended in 500 mL of MBB, applied onto hydrated MS-columns (Miltenyi), washed three times with 500 mL of MBB, and collected with 1 mL of MBB through piston elution.

### High dimensional flow cytometry

SVCs were stained on a polystyrene 96-well V bottom tissue-culture treated plate. The cells were centrifuged at 500 × *g* for 5 min at 4°C, and the cell pellet was washed in PBS once and centrifuged again as described above. The cells were incubated with 25–50 μL of FcBlock (1:100, BD Pharmigen in FACS buffer) on ice for 10 min, and then an equal volume of the staining cocktail was added and mixed (**Table III**). For the identification of the main infiltrated leukocyte populations, total leukocytes were identified gating on CD45^+^ cells. Inside this gate, neutrophils were identified as Ly6G^+^Ly6C^-^ cells and macrophages as CD11b+F4/80+CD64^+^ cells. Ly6C^-^Ly6G^-^ cells were further gated to identify CD3^+^ T-lymphocytes, NK1.1^+^ NK cells and CD19^+^ B-lymphocytes. Samples were acquired on a 13-color Cytoflex (Beckman Coulter) and for each analysis, at least 0.5×10^6^ live cells were acquired by gating on aqua Live/Dead negative cells and upon exclusion of cell doublets (Talamonti et al., 2017;Leuti et al., 2021;Rosina et al., 2022).

**Table III.** Antibodies used for the immunophenotypic characterization.

| **Antibody** | **Source** | **Identifier** |
| --- | --- | --- |
| CD11b APC-Vio770 | Miltenyi Biotec | 130-113-232 |
| CD19 BV421 | Biolegend | 115537 |
| CD3 PE | BD Biosciences | 552127 |
| CD4 PerCP5.5 | Biolegend | 100540 |
| F4/80 FITC | Biolegend | 123107 |
| Ly6C PE-Cy7 | Biolegend | 128018 |
| Ly6G PE-CF594 | BD Biosciences | 15872159 |
| NK1.1 APC | Miltenyi Biotec | 130-120-507 |

### Lactate and nitrite assay

Extracellular lactate was measured in culture medium by an enzyme-based spectrophotometric assay. Cell media were collected and treated with 1:2 (v/v) 30% trichloroacetic acid to precipitate proteins. The resulting mixture was then centrifuged at 14,000×g for 20 min at 4°C, and the supernatant was collected and incubated (30 min, 37°C) with a reaction buffer containing glycine, hydrazine, NAD^+^ and LDH enzyme to allow the conversion of lactate to pyruvate, while simultaneously reducing NAD^+^ to NADH. NADH concentration, stoichiometrically equivalent to the amount of lactate, was determined at 340 nm using a spectrophotometer (extinction coefficient of 6220 M^-1^ cm^-1^). Lactate concentrations were normalized to the total cell protein content. Nitrite concentration was determined by Griess reaction in culture media as previously described (Aquilano et al., 2003).

### Seahorse analysis

Cells were seeded at 1.5 x10^4^ cells/well in a Seahorse XF 96-well plate. Mito-stress test and glycolysis test were performed according to Agilent Technologies recommendations with minor adaptations for adipocytes as previously described (Rosina et al., 2022). Oxygen consumption rate (OCR) and extracellular acidification rate (ECAR) was used as an indicator of mitochondrial function and glycolytic activity, respectively. Analyses were performed using the Desktop Wave Software (Agilent Technologies).

### Targeted metabolomics

Metabolomic data were obtained using liquid chromatography coupled to tandem mass spectrometry. We used an API-3500 triple quadrupole mass spectrometer (AB Sciex, Framingham, MA, USA) coupled with an ExionLC™ AC System (AB Sciex). Cells were disrupted in a tissue lyser for 1 min at maximum speed in 250 µl ice-cold methanol:acetonitrile 1:1 (v/v) containing 1 ng/µl [U-^13^C_6_]-glucose and 1 ng/µl [U-^13^C_5_]-glutamine as internal standards. Lysates were spun at 15,000 g for 15 min at 4°C. Samples were then dried under N_2_ flow at 40°C and resuspended in 5 mM ammonium acetate in methanol:water 1:1 (v/v) for subsequent analyses.

As previously described, quantification of amino acids, their derivatives, and biogenic amines was performed through previous derivatization (Audano et al., 2018). Briefly, 25 µl of each 125 µl sample were collected and dried separately under N_2_ flow at 40°C. Dried samples were resuspended in 50 µl phenyl-isothiocyanate, EtOH, pyridine, and water 5%:31.5%:31.5%:31.5%, then incubated for 20 min at RT, dried under N_2_ flow at 40°C for 90 min, and finally resuspended in 100 µl 5 mM ammonium acetate in MeOH/H_2_O 50:50. Quantification of different amino acids was performed using a C18 column (Biocrates, Innsbruck, Austria) maintained at 50°C. The mobile phases for positive ion mode analysis were phase A: 0.2% formic acid in water and phase B: 0.2% formic acid in acetonitrile. The gradient was T_0_: 100%A, T_5.5_: 5%A, T_7_: 100%A with a flow rate of 500 µl/min. All metabolites analysed in the described protocols were previously validated by pure standards and internal standards were used to check instrument sensitivity.

Quantification of energy metabolites and cofactors was performed using a cyano-phase LUNA column (50mm x 4.6mm, 5 µm; Phenomenex) with a 5.5 min run in negative ion mode with two separated runs. Protocol A: mobile phase A was water and phase B was 2 mM ammonium acetate in MeOH, with a gradient of 10% A and 90% B for all analyses and a flow rate of 500 µl/min. Protocol B: mobile phase A was water and phase B was 2 mM ammonium acetate in MeOH, with a gradient of 50% A and 50% B for all analyses and a flow rate of 500 µl/min.

Acylcarnitine quantification was performed on the same samples using a Varian Pursuit XRs Ultra 2.8 Diphenyl column (Agilent). Samples were analysed in a 9 min run in positive ion mode. Mobile phases were A: 0.1% formic acid in H_2_O, B: 0.1% formic acid in MeOH, and the gradient was T_0_: 35%A, T_2.0_: 35%A, T_5.0_: 5%A, T_5.5_: 5%A, T_5.51_: 35%A, T_9.0_: 35%A with a flow rate of 300 µl/min. MultiQuant™ software (version 3.0.3, AB Sciex) was used for data analysis and peak review of chromatograms. Raw areas were normalized to the areas’ median. Obtained data were then compared to controls and expressed as fold change. Raw data are reported in **Suppl. Table I**.

### 16s Sequencing analysis

Upon collection, faecal pellets were immediately preserved by freezing in liquid nitrogen and then stored at −80°C. To extract faecal nucleic acid, an E.Z.N.A. stool DNA kit (OMEGA, Bio-tek) was used. The bacterial 16S rRNA gene was amplified from total DNA following the Illumina 16S Metagenomic Sequencing Library Preparation instructions, targeting the V3-V4 hypervariable region amplicon by PCR with universal primers containing Illumina adapters reported in Klindworth et al. (Klindworth et al., 2013). The resulting amplicon was purified and subjected to a second PCR to barcode the libraries using the Illumina dual-index system before a final purification step. The pooled libraries were then sequenced using paired-end sequencing (2×300 cycles) on an Illumina MiSeq device according to the manufacturer’s specifications. The resulting sequence data obtained as FASTq files were analysed using 16S Metagenomics GAIA 2.0 software, which performs quality control on the reads/pairs (i.e., trimming, clipping, and adapter removal) through FastQC and BBDuk before mapping them with BWA-MEM against NCBI databases. The average number of reads per sample was 220,507.1 (SD ± 104,675.1).

### Statistical analysis

Data were expressed as the mean ± SD. The exact numbers of replicates are given in each figure legend. A two-tailed unpaired Student’s *t*-test was performed to assess the statistical significance between two groups. One-way analysis of variance followed by Dunnett’s (comparisons relative to controls), or Tukey’s (multiple comparisons among groups) *post hoc* tests was used to compare three or more groups. Statistical analyses were performed using GraphPad Prism 9 (GraphPad Software Inc., San Diego, CA, USA). In all cases, a *p*-value of 0.05 was set as the significance threshold.

## Acknowledgements

This work was mainly supported by Friedreich’s Ataxia Research Alliance (FARA) - General Research Grant 2020 to K.A. Other supports: FARA - General Research Grant 2021 to D.L.-B.; Progetto Eccellenza 2018–2022 to the Dipartimento di Scienze Farmacologiche e Biomolecolari, Università degli Studi di Milano to S.P. and N.M., and the Italian Ministry of Health (Ricerca Corrente and 5 × 1,000 funds) to N.M. S.P. is supported by Fondazione Umberto Veronesi postdoctoral fellowship.

## Authors’ contributions

R.T. and F.S. carried out the experiments, sample and data collection. M.T. and V.C. performed cytofluorimetric analyses; M.A., S.P. performed metabolomic analyses; C.P. and V.P. performed metagenomic analyses, V.N. carried out Seahorse analyses; S.C. carried out histochemical analyses; N.M. contributed essential reagents. D.L.-B contributed to the original idea, helped supervision and analysed the data. K.A. conceived the original idea, acquired funding, wrote the manuscript and supervised the experiments. V.P., A.F., S.C., V.C., N.M. and D.L.-B contributed to the final version of the manuscript.

## Data availability

Fastq files of 16s metagenomic analysis are deposited in ArrayExpress (E-MTAB-12871). Bulk RNAseq data are available through Gene Expression Omnibus gene repository with the dataset identifier *in progress*. Any additional raw data is available from the lead contact upon reasonable request.

